# Multiplexed single cell transcriptomics optimizes mesodermal patterning and hemogenic endothelial output from murine embryonic stem cells

**DOI:** 10.1101/2024.05.24.595249

**Authors:** Barbara Varnum-Finney, Adam M. Heck, Sanjay R. Srivatsan, Stacey Dozono, Rachel Wellington, Alisa B. Chankhunthod, Cynthia Nourigat-McKay, Tessa Dignum, Cole Trapnell, Brandon Hadland

## Abstract

**Background:** Early patterning of mesodermal precursor populations is a key step of hematopoietic development in the embryo. To better understand this process, we employed sci-Plex, a high-throughput method of measuring multiplexed perturbations at the single-cell level, to evaluate the transcriptional response of mouse embryonic stem cells subjected to a gradient of two key morphogens in early mesoderm/hematopoietic development, Activin and BMP4.

**Results:** sci-Plex revealed varying combinations of Activin and BMP4 temporally influenced mesoderm patterning in vitro and subsequent production of cell types reflecting their in vivo counterparts. We leveraged sci-Plex data to further optimize the generation of intraembryonic-like hemogenic endothelial cells that serve as the precursors of definitive hematopoietic lineages, including hematopoietic stem cells.

**Conclusions:** This study highlights the utility of sci-Plex to dissect how dose and temporal integration of interacting signal pathways determines cell fates and serves as a resource to analyze cell fate choices in early mesoderm patterning at single cell resolution.

## Background

Hematopoietic stem cells (HSCs) are clinically valuable due to their distinctive capacity for life-long reconstitution of all hematopoietic lineages in the marrow of conditioned transplant recipients. Yet, it has proven particularly difficult to reproducibly derive long-term engrafting HSCs from either murine embryonic stem cells (mESC) or human induced pluripotent stem cells (huiPSC) despite decades of efforts (1). This is primarily due to the complexity of embryonic hematopoiesis such that attempts at recapitulating HSC genesis in ESC culture have resulted in a multiplicity of false leads (2). Embryonic hematopoiesis occurs in temporal waves of different hematopoietic progenitor types prior to the emergence of the first HSCs (3). The initial wave, referred to as the primitive wave, occurs almost immediately after gastrulation in extraembryonic tissues, generating the first erythroid cells and megakaryocytes (4, 5). Subsequent waves of hematopoiesis are derived from specialized endothelial cells known as hemogenic endothelium (HE), giving rise to blood cell fates through a process referred to as the endothelial to hematopoietic transition (EHT). EHT occurs in various embryonic sites including the yolk sac, placenta, endocardium, and the aorta-gonad-mesonephros region (AGM), first generating HSC-independent oligopotent and multipotent progenitor types with various erythroid, myeloid and lymphoid potential, followed by the emergence of the first long-term engrafting HSCs (6–11). Many mESC and huiPSC hematopoietic differentiation protocols have recapitulated the HSC-independent progenitor waves in vitro. However, most protocols developed to date have failed to robustly and reproducibly generate HSCs. Recently reports provide important proof of principle for the generation of HSC-like populations from huiPSC capable of multilineage engraftment in immune-deficient mouse models (12, 13). However, the production of transplantable huiPSC-derived HSCs remains highly inefficient, and their serial engraftment capacity remains inferior to that of developmentally matched HSCs derived from the human embryo (14). This highlights the need to better understand the sequential embryonic cell types generated in current differentiation protocols in vitro and how they relate to the populations in the embryo that serve as precursors to the successive waves of hematopoiesis in vivo, including HSCs.

Mouse gastrulation starts with primitive streak (PS) formation at the extra-/intra-embryonic boundary around embryonic day 6-6.5. The position of PS formation defines the posterior pole of the embryo, and expression of the T-box transcription factor Brachyury is an early indicator of PS formation. As the PS elongates from this posterior position anteriorly, cells continuously exit and migrate from the elongating PS (15). Those migrating anteriorly have an embryonic fate (intra-embryonic cells (IEC)) while those migrating proximally have an extra-embryonic fate (extra-embryonic cells (EEC)) (16). Following departure from the PS, IEC migrate toward the embryo proper, forming the bilateral wings of embryonic mesoderm that eventually gives rise to paraxial mesoderm, pharyngeal mesoderm, and other embryonic tissues as well as lateral plate mesoderm (LPM), that produces cardiac cells and contributes to multilineage hematopoiesis including the first HSCs (17). EECs migrate toward the extra-embryonic area and give rise to extra-embryonic tissues including the hematopoietic cells of the first primitive waves as well as the extra-embryonic mesoderm that gives rise to a subsequent yolk sac wave of HE-derived erythromyeloid progenitors (EMP). Morphogens, such as BMP4, Wnt and Activin/Nodal, established in concentration gradients within the embryo, direct these migrations and further promote differentiation to specific cell types that differentially contribute to intraembryonic and extraembryonic tissues. It is well established that mesoderm differentiation in mESC and huiPSC in vitro culture also requires these morphogens (BMP4, Wnt, and Activin/Nodal) in a time and concentration-dependent manner (18–21). Protocols for in vitro differentiation of mESC and huiPSC have not been able to fully assess the outcome of morphogen-dependent gradients on mesodermal/hematopoietic differentiation due to a lack of available high-throughput tools to characterize cell fate responses to multiplexed combinations of morphogen concentrations at the single cell level. To overcome this barrier, we applied sci-Plex, a novel method to measure transcriptional responses to highly multiplexed perturbations at single-cell resolution (22), to examine the combinatorial dose and time-dependent effects of exogenous Activin and BMP4 gradients on mesodermal patterning and hematopoietic differentiation from mESC. This approach identified mESC-derived cell fates responsive to combinatorial Activin and BMP4 gradients that recapitulate those emerging in both the intra-embryonic and extra-embryonic regions during early murine embryogenesis. Furthermore, the technique facilitated optimization of a differentiation protocol to enhance LPM and HE output as an essential initial step towards HSC production from pluripotent stem cells in vitro.

## Methods

### Mice

Wild-type C57Bl6/J7 (CD45.2) and congenic C57BL/6.SJL-Ly5.1-Pep3b (CD45.1) mice were bred at the Fred Hutchinson Cancer Research Center. Male and female C57Bl6/J7 CD45.2 mice at 8–12 weeks of age were used for transplantation experiments. All animal studies were conducted in accordance with the NIH guidelines for humane treatment of animals and were approved by the Institutional Animal Care and Use Committee at the Fred Hutchinson Cancer Center.

### Mouse embryonic stem cells (mESCs)

mESCs from frozen aliquots of the same derivation were used for all differentiation experiments at less than passage 10. Briefly, cells were thawed fresh for each differentiation experiment and cultured in gelatinized, tissue culture 6-well plates or 60 mm dishes in ESGRO Complete PLUS Clonal Grad Medium (MilliporeSigma). Cells were passaged every other day using TrypLE to dissociate into a single cell suspension for 2-4 passages before being seeded for embryoid body differentiation

### AGM-derived Akt-ECs (AGM-ECs)

AGM-EC were generated as previously described (23) and further detailed in a protocol available at Nature Protocol Exchange (https://protocolexchange.researchsquare.com/). For reproducibility, AGM-EC from frozen aliquots of the same derivation were used for all co-culture experiments at less than passage 15 for experiments in this manuscript.

AGM-EC were plated at a density of 5×105 cells/flask in gelatinized, tissue culture T75 flasks in EC media (consisting of IMDM with 20% FBS, Penicillin/streptoMycin, Heparin, L-glutamine, and EC mitogen), and passaged every 3–4 days when confluent.

### OP9/OP9-DLL4 stromal cells (OP9 cells)

GFP-expressing OP9 bone marrow stromal cells and GFP-expressing OP9 bone marrow stromal cells ectopically expressing Delta-like 4 (OP9-DLL4 cells) generated by the Zúñiga-Pflücker lab (24) were used to assay the B- and T-lymphoid potential of hematopoietic colony cells. OP9 cell lines were plated at a density of 1.6×105 cells/flask in gelatinized, tissue culture T25 flasks in OP9 media (consisting of aMEM with 20% FBS, and Penicillin/streptoMycin). Cells were passaged every 3 days (or when confluent) and used at passage numbers below 13 for co-culture assays

### Embryoid body differentiation and cell sorting

mESCs were collected and dissociated into single cell suspension by incubation in Accutase. Cells were washed in serum-free media, resuspended in StemPro-34 serum-free media and plated onto 100 mm petri dishes at a density of 2×10^6^ cells per plate to allow for EB formation. EBs were cultured in StemPro-34 for 51 hours, EBs were then collected into 15 mL conical tubes (no more than 2x 100 mm dishes per tube) and allowed to gravity settle for 10 minutes at 37°C. Once settled, media was gently removed, 1 mL TrypLE per dish was added, and cells were resuspended by gently flicking the tube 2-3 times. Cells were incubated in 37°C water bath for 3 minutes with gentle swirling and 1-2 flicks halfway through the incubation to keep the EBs suspended. An equivalent volume of StemPro-34 was added and cells were gently triterated 10x times with a 1000 uL pipette to fully dissociate EBs. Cells were then pooled, if applicable, washed in serum-free media, and resuspended in 300 uL of StemPro-34. 30 uL of DNase was added and cells were incubated for 10 minutes at room temperature. Following DNase treatment, cells were washed in serum-free median and resuspended in StemPro-34 containing recombinant vascular endothelial growth factor (VEGF) at 10 ng/mL and a combinatorial gradient of Activin and bone morphogenetic protein 4 (BMP4). It was determined a combination of 20 ng/mL Activin and 5 ng/mL BMP4 yielded the most robust downstream hematopoietic output, and was used in subsequent experiments unless otherwise noted. Cells were plated onto 24-well low-adherence plates at 4×10^5^ cells per well to allow for EBs reformation. Following 72 hours of incubation (Day 5 of the differentiation protocol), the Activin concentration in the media was lowered. To do this, EBs were collected, gravity settled, and media was replaced with StemPro-34 containing 10 ng/mL VEGF, 5 ng/mL Activin and BMP4. When noted in certain experiments, all-trans retinoic acid (ATRA) was added at this point as well. Following media swap, EBs were gently replated onto new 24-well low-adherence plates.

After another 24-48 hours of culture (Day 6 or 7 of differentiation), dependent on the experiment, cells were sorted to isolate hematopoietic precursors and/or endothelium. Briefly, EBs were collected and dissociated to single-cell suspension as described above. Cells were then washed with PBS containing 10% FBS. Cells were incubated with anti–mouse CD16/CD32 (FcγRII block) and stained with the following monoclonal antibodies as indicated. Relevant isotype control antibodies were used to set gates. DAPI staining was used to gate out dead cells. All reagents for cell staining were diluted in PBS with FBS, and staining was carried out on ice or at 4°C. Cells were sorted on a BD FACS Aria II. Sorted cells were subjected to downstream co-culture, lineage or transplantation assays as described below.

As noted for Sci-Plex experiments, EBs were collected and dissociated to single-cell suspension as described above over the course of differentiation (day 4, 5, and 6) to assess cell surface phenotype by flow cytometry or transcriptional expression by Sci-plex. Dissociated cells were processed for Sci-plex library prep and sequencing as previously described (22).

### Flow cytometry analysis

At various timepoints during differentiation or following co-culture, a fraction of the cell population (approximately 10-20% unless otherwise stated) was harvested for analysis of surface phenotype by flow cytometry. For differentiation timepoints, EBs were dissociated by incubation in TrypLE. Following co-culture, hematopoietic progeny were harvested from the EC layer by vigorous pipetting. Following collection, cells were spun and re-suspended in PBS with 2% FBS, pre-incubated with anti-mouse CD16/CD32 (FcRII block) and then stained with anti-mouse monoclonal antibodies as indicated. DAPI was used to exclude dead cells. Flow cytometry was performed on a Becton Dickinson Canto 2 and data analyzed using FlowJo Software.

### AGM-EC co-culture

Co-culture experiments were carried out with AGM-EC at passage 15 or less plated 24-48 hours prior to initiation of co-culture onto gelatin-treated 96-well or 24-well tissue culture plates at a density of 1×10^4^ or 5×10^4^ cells per well, respectively. Sorted mESCs were re-suspended in serum-free X-vivo 20 culture media with recombinant cytokines: stem cell factor (SCF) and FMS-like tyrosine kinase 3 ligand (FLT3L) each at 100 ng/ml, and interleukin-3 (IL-3) and thrombopoietin (TPO) each at 20 ng/ml. Following 5 to 7 days of co-culture hematopoietic progeny were harvested by vigorous pipetting for subsequent analysis by flow cytometry and/or lineage assays, as described below.

### Colony forming unit (CFU) assays

In preparation for CFU assays, methylcellulose-based medium with recombinant cytokines was plated to 96-well non-tissue culture plates. Hematopoietic progeny were harvested by vigorous pipetting from the EC layer, washed in serum-free media, and resuspended in serum-free media. 10% of the cells from each well/replicate were added to individual 96-wells containing methylcellulose/cytokines (the remaining 90% was used for OP9 co-culture as described below). Visual characterization of resultant CFU colonies—CFU-Erythroid (CFU-E), CFU-granulocyte/macrophage (CFU-GM), or multilineage colonies (CFU-GEMM) was performed under a light microscope after 10– 12 days of culture.

### OP9 co-culture assays

OP9 and OP9 DLL4-expressing (OP9-DLL4) cells were plated 24–48 hours prior to OP9 co-culture initiation at a density of 1×104 cells/well to gelatin-treated 24-well tissue culture plates in OP9 media. On the day of co-culture, 1mL/well of OP9 co-culture media containing FLT3L and interleukin-7 (IL-7) at 5 ng/ml each was added to OP9 24-wells in place of the OP9 media used for plating. The remaining 90% of hematopoietic co-culture cells processed as described above were divided equally and plated to OP9 or OP9-DLL4 coated wells.

After 10–12 days of co-culture, 50% of cells from the OP9 wells were harvested for flow cytometry analysis. Cells were stained in PBS with 2% FBS containing anti-mouse CD16/32 (FcRII block) and monoclonal antibodies AA4.1 (PE), B220 (Per-CP), CD19 (APC), and CD11b (APC-eFluor780). DAPI was used to exclude dead cells and contaminating GFP-expressing OP9 cells were excluded by gating on the FITC-negative population. Flow cytometry was performed on a Becton Dickinson Canto 2 and data analyzed using FlowJo Software. Isotype control-stained cells were used to set gates for analysis. B-lymphoid cells were identified phenotypically as CD11b−AA4.1+CD19+B220+.

Every 5–8 days of co-culture (or when cells began to overgrow), 50% of cells from OP9-DLL4 co-culture were transferred to newly plated OP9-DLL4 wells. After 12–21 days total, 50% of cells from OP9-DLL4 wells were harvested for flow cytometry analysis. Cells were stained in PBS with 2% FBS containing anti-mouse CD16/32 (FcRII block) and monoclonal antibodies CD25 (PE-Cy7), CD44 (APC), CD4 (PerCP), and CD8 (PE). DAPI was used to exclude dead cells, and contaminating GFP-expressing OP9 stromal cells were excluded by gating on the FITC-negative population. Flow cytometry was performed on a Becton Dickinson Canto 2 and data analyzed using FlowJo Software. Isotype control-stained cells were used to set gates for analysis. T cells were identified phenotypically as CD25+CD44−CD4+CD8+.

### Transplantation assays

Following co-culture, a fraction of the hematopoietic cells generated were harvested from the EC layer by vigorous pipetting, washed with 2% FBS and re-suspended in 100 uL PBS/2% FBS per mouse transplanted. Samples were supplemented with 5×10^4^ whole marrow cells from adult congenic C57BL/6.SJL-Ly5.1-Pep3b (CD45.1) mice in 100 ul PBS/2% FBS to provide hematopoietic rescue. Cell suspensions (200 ul total volume/mouse) were injected into lethally irradiated (1,000 cGy using a Cesium source) congenic CD45.1 adult recipients via the tail vein. Flow cytometry analysis of peripheral blood obtained by retro-orbital bleeds was performed at various intervals between 2 and 24 weeks following transplantation. Lineage-specific staining for donor (CD45.2) and recipient/rescue (CD45.1) cells from peripheral blood was performed as previously described (23), using anti-mouse monoclonal antibodies indicated: CD45.1, CD45.2, CD3, CD19, Gr1, and F4/80.

### Embryo dissections for Sci-Plex

Embryos were harvested from pregnant females, as previously described (23). Embryo age was determined by morphology (E6.5-7) or by counting somite pairs (E8.5-9). For E6.5-7 and E8.5-9, whole embryo/extra-embryonic tissue, or yolk sac and P-Sp region of the embryo proper were collected, respectively. Dissected tissues were treated with 0.25% collagenase for 30 minutes at 37°C, pipetted to single cell suspension, and washed with PBS containing 10% FBS. The resulting single cell suspension was process for Sci-plex library prep and sequencing as previously described (22).

### Sci-Plex data and analysis

All raw and processed sequencing data is publicly available through GEO (accession number GSE343689). All scripts for single cell transcriptomic analysis will be publicly available on the FHCC GitHub at https://github.com/FredHutch/mESC-SciPlex.

### Dimensionality reduction, batch correction, and cluster analysis

sci-Plex data was analyzed using Monocle3 (25). The preprocess_cds() function was used to project the data onto the top principal components. EB and Embryo datasets were analyzed separately with the num_dim set to 98 and 50, respectively. Uniform Manifold Approximation (UMAP) was used for dimensionality reduction with the reduce_dimension() function, with umap.n_neighbors = 4L. set.seed was used to specify the number of seeds to avoid variability in output due to a random number generator in the function. Clustering was performed with the cluster_cells function with a resolution of 3.6e-4 and 3e-4 for the EB and Embryo datasets, respectively.

### Cell type classification

Cell type classification was performed using diagnostic marker genes curated from Puijuan-Sala et al. 2019 (26) (marker genes listed in Supplementary Table 1). Cell clusters were classified as a given cell type based aggregate expression of diagnostic markers (see heatmap in Figure S4A). Using this method, a small population of cells remains unclassified rather than inputting cell types for all cells in the scRNA-seq dataset. Cell type classifications were used for all downstream analyses, as indicated.

### Gene-set scores

Gene set scores were calculated for each single cell as the log-transformed sum of the size factor-normalized expression for each gene in published signature gene sets as follows; primitive streak (26), extra-embrynoic (EEC) and intra-embryonic (IEC) (16), lateral plate mesoderm (LPM) (27, 28), and arterial endothelial cells (29, 30). A list of the genes and associated references can be found in Table 2.

### Sub-setting and cell counting

Cells were subset or groups of cells were highlighted based on cell classification and/or specific gene expression as labeled. Cell counts for a given group were generated from cell dataset metadata for Activin/BMP4 concentration and/or collection timepoint. All heatmaps and cell dot plots were generated in the ggplot2 package.

### Quantification and statistical analysis

Statistical analyses of colony frequencies were conducted with Prism. Data are expressed as mean ± standard deviation (s.d.), and n indicates biological repeats, unless indicated otherwise. P values calculated by Student’s t test unless otherwise indicated.

## Results

### sci-Plex reveals early embryonic cell fates resulting from exposure of differentiating mESC to combinatorial doses of Activin and BMP4

In order to control for exogenous factors, we built upon a serum-free protocol for mESC differentiation that was previously shown to generate mesoderm capable of producing multilineage hematopoietic cells (31, 32). To mimic Activin/Nodal and BMP4 gradients that pattern the embryo during PS formation, we applied 16 different combinations of Activin and BMP4 concentrations (19, 21) and used sci-Plex to trace mesodermal differentiation and subsequent cell fates at single cell resolution at various time points in the differentiation protocol (Figure 1A). Briefly, mESCs were cultured as embryoid bodies (EBs) for two days without exogenous growth factors, since previous studies have shown that mESC are unresponsive to Activin or BMP4 for 24 to 48 hours following removal of LIF or growth media (18). At 48 hours, EBs were dissociated to single cells and re-suspended to form EBs in a 4×4 gradient of Activin and BMP4, along with a consistent concentration of VEGF required for hematovascular fates (33). On days 4, 5 and 6 of culture (days 2, 3 and 4 following addition of growth factors), EBs were dissociated, and a portion of each sample was processed for both sci-Plex and analyzed for immunophenotypic markers by flow cytometry. As expected, the immunophenotypic data indicates a range of mesodermal and endothelial outputs responsive to different Activin and BMP4 concentrations over the period of differentiation analyzed, including Pdgfra^+^Flk1^+^ LPM-like cells previously shown to generate both cardiac and hematovascular lineages (20, 34) (Figure S1A) and Flk^+^VE-Cadherin^+^ cells indicating endothelial differentiation (Figure S1B).

**Figure 1:**
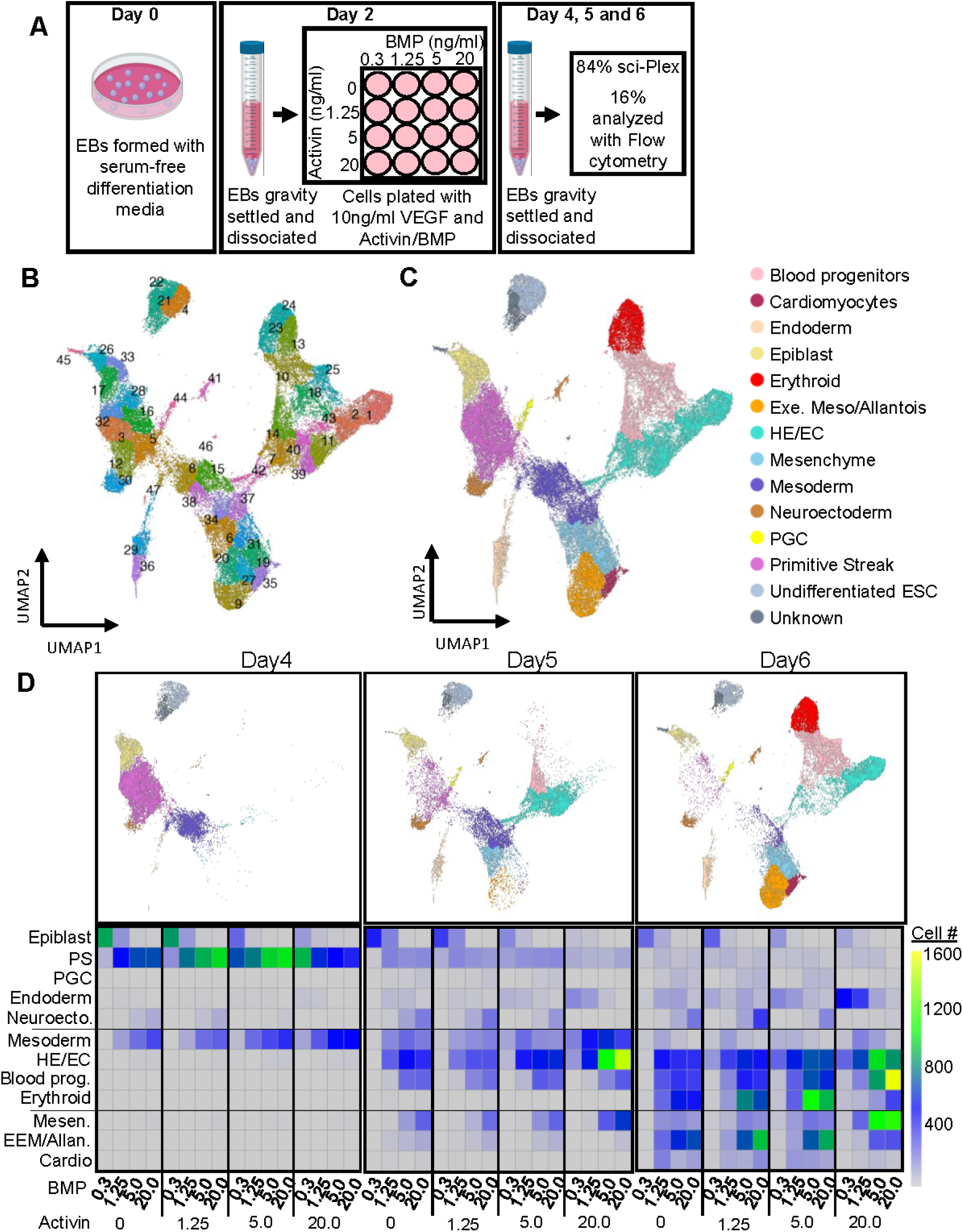
sci-Plex analysis reveals cell types generated in embryoid body differentiation. A: Diagram of sci-Plex experimental setup. B: UMAP and cluster analysis of scRNA-seq data from EB differentiation sci-Plex experiment. C: Cell type classification based on marker genes (see methods). D: (Top) Cell type classification for each differentiation time point collected. (Bottom) Heatmaps for number of cells generated for notable cell types per condition across differentiation time points collected.

sci-Plex data was analyzed using Monocle3 (25) by first performing dimensionality reduction by uniform manifold approximation and projection (UMAP) and unsupervised cluster analysis of the single-cell transcriptomes (Figure 1B). Gene expression levels for previously published diagnostic marker genes from early mouse embryonic development were used to assign cell types to each cluster (Figure 1C; Table 1; Figure S2A) (26). As expected, analysis across time points revealed a clear progression from PS to mesoderm to differentiated mesoderm-derived cell types (Figure 1D). Moreover, different transcriptionally defined cell types were observed across the spectrum of Activin and BMP4 concentrations (Figure S2B) and depiction of the relative cell number by day and concentration demonstrated concentration-dependent effects on the generation of each cell type (Figure 1D). Mesoderm was maximally generated on days 4 and 5 at the highest Activin/BMP4 concentrations. Endothelial cell types (HE/EC) were detected on days 5 and 6 at all Activin concentrations but were maximally generated at the highest Activin and BMP4 concentrations. Differentiating blood progenitors and erythroid cells were maximally generated on day 6, particularly at higher BMP4 concentrations, whereas the highest Activin concentration inhibited generation of erythroid cell types. These results are consistent with previous studies that have shown hematopoiesis occurs in waves within the differentiating EB, similar to waves of hematopoiesis seen in the embryo (32). This first wave resembles the yolk sac primitive hematopoiesis, generating erythroid cells predominantly expressing the characteristic Beta-H1 hemoglobin (*Hbb-bh1*) (Figure S2C). Our observation that a primitive erythroid-like wave is maximally generated at higher BMP4 and lower Activin is also consistent with the embryonic axis where higher BMP4 is found posteriorly associated with yolk sac mesoderm and higher nodal/activin more anteriorly with intra-embryonic mesoderm formation. We found cells expressing cardiac markers predominately at lower concentration of BMP4 (Figure 1D and S2B), consistent with previous publications (21). Higher Activin levels were required for endoderm generation whereas higher BMP4 was inhibitory, consistent with distal endoderm formation in the embryo (35) and endoderm differentiation in huiPSC studies (36). Measurable numbers of primitive germ cells (PGC) were only seen at higher BMP4 concentrations and were inhibited by the highest Activin concentration, consistent with previous studies showing BMP4 is required for PGC formation in a murine organ culture system (37). Together, these studies demonstrate that sci-Plex robustly captures transcriptionally defined cell fates during mESC differentiation in vitro, identifying both temporal relationships and concentration-dependent effects on cell fates that recapitulate the expected developmental trajectories of their embryonic counterparts in vivo.

### Intra-embryonic and extra-embryonic mesodermal fates are patterned by Activin and BMP4

In the embryo, cells within the primitive streak express T-box transcription factor *Brachyury*, but downregulate its expression as they migrate within the mesoderm wings and differentiate into mesoderm (38). During mESC differentiation, cells expressing *Brachyury* were identified predominantly on day 4, two days following the addition of BMP4, Activin and VEGF (Figure 2A and B), consistent with previous studies (32). As expected, *Brachyury* expression was downregulated as EB cells increasingly expressed mesodermal markers *Pdgfra* and *Kdr* between day 4 and 6, which was accelerated at higher Activin concentrations. EBs exposed to the highest Activin concentration contained the most cells expressing both *Flk1* and *Pdgfra* on both days 4 and 5 (Figures 2A and B), demonstrating that the pattern of gene expression by day and concentration revealed by sci-Plex is remarkably similar to that observed based on protein expression by flow cytometric analysis (compare Figures S1A and 2B).

**Figure 2:**
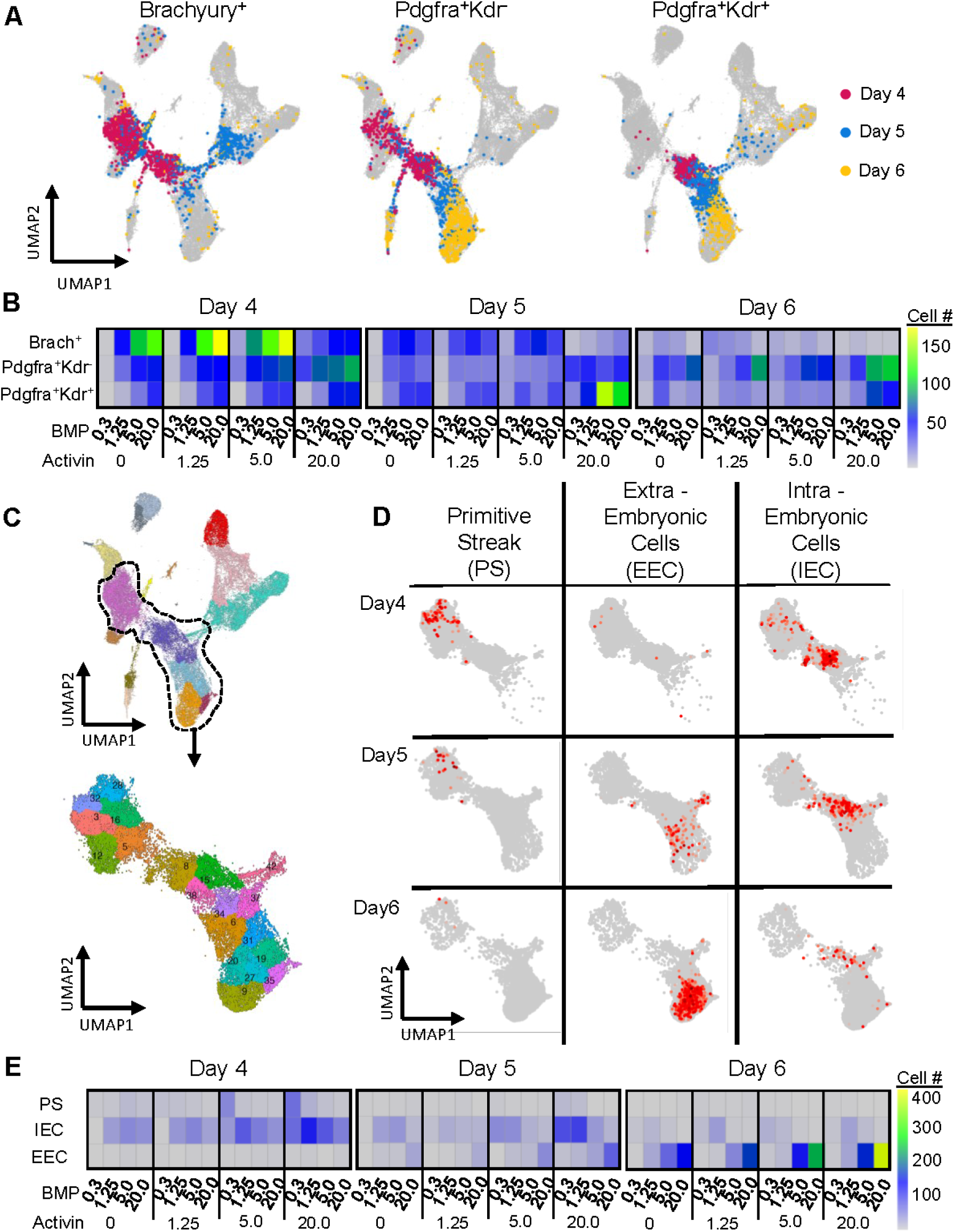
Activin and BMP4 exposure dictates early cell fate patterning. A: Gene expression of early mesodermal cell fate markers across differentiation time points collected. B: Heatmaps for number of cells generated per cell fate markers from (A), across differentiation time points collected. C: Subset of clusters representing primitive streak and mesodermal patterning in UMAP space. D: Heatmaps of gene-set scores for signature gene sets defining primitive streak (26), extra-embryonic cells and intra-embryonic cells (16) within the primitive streak and mesodermal subset from (C). E: Heatmaps for number of cells above the gene set midpoint from (D) across differentiation time points collected.

Embryonic day E7.5-8 EECs exiting the PS express genes associated with hematopoiesis such as *Tal1* and *Runx1*, consistent with their rapid differentiation to produce the first primitive hematopoietic wave, whereas exiting IECs express genes generally associated with mesoderm (16). To map and determine numbers of cells bearing genes characteristic of PS, IEC and EEC, we determined relative aggregate expression gene-set scores (GSS) within the subset of EB clusters representing PS and early mesodermal patterning (Figure 2C). Gene-sets were generated from previous publications (Table 2). Cells with high PS GSS were primarily observed on day 4, while cells with the highest IEC or EEC GSS were found in a generally non-overlapping pattern, as expected (Figure 2D). Most cells with a high IEC GSS at all concentrations were observed on days 4 and 5, whereas those with highest EEC GSS were observed predominantly on days 5 and 6 (Figure 2E). Higher Activin generated more cells with high IEC GSS on days 4 and 5, whereas higher BMP4 inhibited generation of IEC and promoted generation of EECs on days 5 and 6 (Figure 2E). These results are consistent with the observation that EECs exit from the posterior PS where higher BMP4 levels are present, whereas IECs exit more anteriorly at higher Activin/Nodal concentrations (16).

### sci-Plex reveals optimized conditions for generation of LPM and HE

In the embryo, LPM is the source of HE that generates intra-embryonic multilineage hematopoietic progenitors and HSCs (17). Cells with high LPM GSS were observed by day 4, peaked at day 5, and were diminished by day 6 with progression to hematovascular or other mesodermal fates, including cardiac (Figures 3A). Higher Activin concentrations promoted LPM compared to lower Activin concentrations, even at the lowest BMP4 concentration, with optimal LPM generation occurring at day 5 in conditions with the highest concentration of Activin (20ng/ml) and intermediate BMP4 (5ng/ml) (Figure 3A).

**Figure 3:**
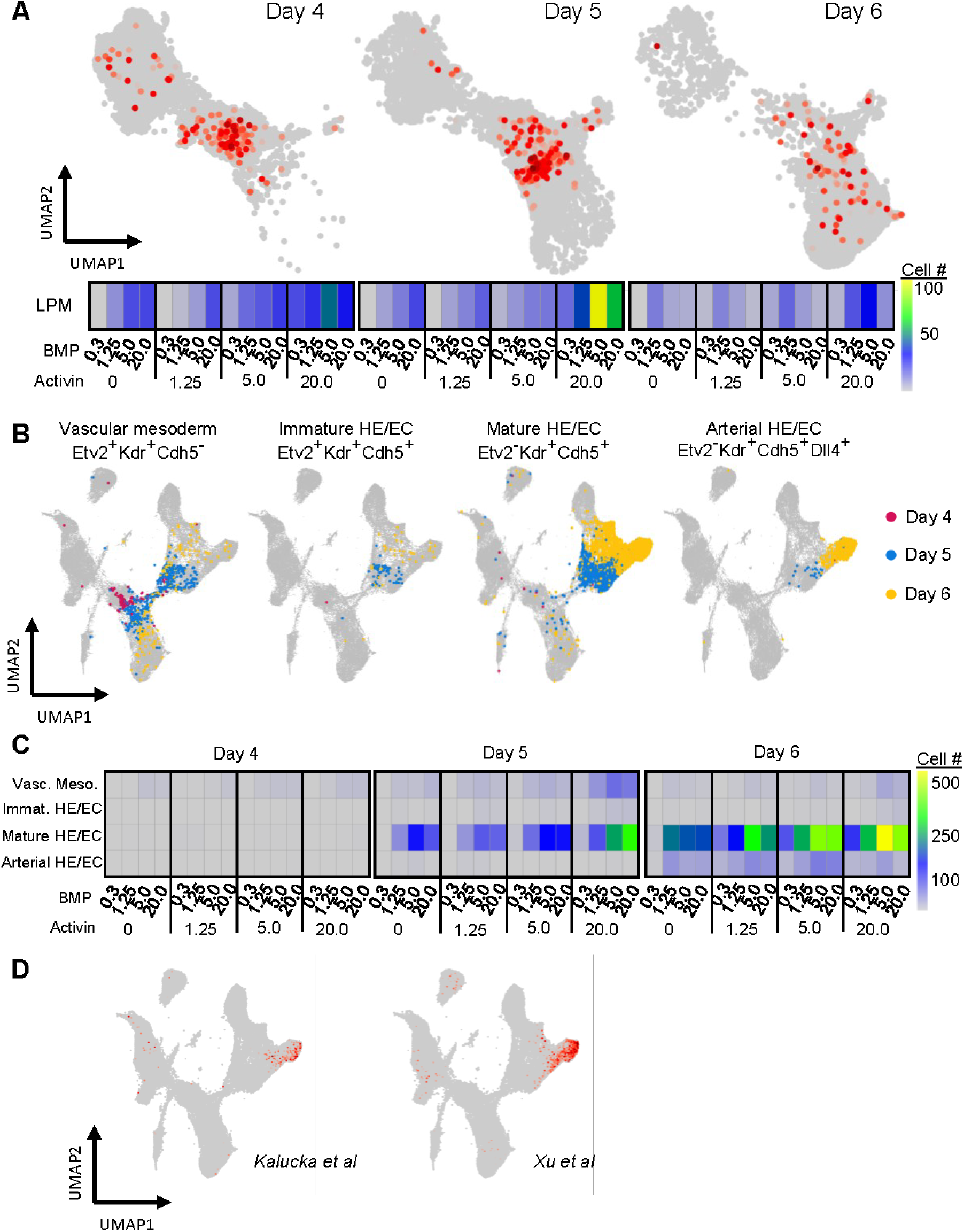
sci-Plex reveals optimal Activin/BMP conditions for generation of LPM and HE. A: (Top) Heatmap for gene-set scores for lateral plate mesoderm signature gene set (27, 28) within the primitive streak and mesodermal subset from (2C). (Bottom) Heatmaps for number of cells above the gene set midpoint across differentiation time points collected. B: Gene expression of hematovascular cell fate markers across differentiation time points collected. C: Heatmaps for number of cells generated per cell fate markers from (B), across differentiation time points collected. D: Heatmaps of gene-set scores for signature gene sets defining arterial endothelial cells (29, 30), across the entire sci-Plex dataset.

In the embryo proper, LPM (co-expressing *Flk1* and *Pdgfra*) destined for hematovascular fates simultaneously downregulate *Pdgfra* and increase expression of *Etv2*, a gene essential for generating vascular and hemogenic endothelium (34, 39). Subsequently, a subset of *Flk1*^+^*Etv2*^+^ mesoderm cells upregulate VE-Cadherin (*Cdh5*) as they mature to differentiated endothelial cell types (HE/EC). Consistent with this developmental progression, our sci-Plex data revealed substantial numbers of *Flk1^+^Etv2^+^Cdh5*^-^ hematovascular mesoderm and *Flk1^+^Etv2^+^Cdh5^+^* immature HE/EC predominantly on day 5 (Figures 3B and C). Similar to LPM, hematovascular mesoderm and immature HE/EC were optimally generated at 20ng/ml Activin and 5ng/ml BMP4. Downregulation of *Etv2* is required for further EC/HE maturation prior to the emergence of hematopoietic and various EC (arterial, venous, lymphatic) fates. Consistent with this, *Kdr^+^Etv2^-^Cdh5^+^* cells are increased between days 5 and 6 in our EB sci-Plex data, and by day 6, a subset also express *Dll4*, a Notch ligand expressed in arterial EC and HE (Figure 3B) (40, 41). To validate the emergence of arterial HE/EC in the EB sci-Plex data, we calculated arterial endothelium GSS using two sets of arterial endothelial markers (Figure 3D) (29, 30). In contrast to immature HE/EC, the highest Activin concentration generated fewer arterial HE/EC than lower activin concentrations, suggesting that persistent Activin exposure may be inhibitory to generation of arterial EC/HE from immature HE/EC (Figure 3C). Based on this result, we revised our protocol to shift EBs to a lower Activin concentration on day 5 to enhance arterial EC/HE production.

### EBs contain both yolk sac-like and intraembryonic-like HE

Yolk sac HE have been show to express LYVE1 (lymphatic vessel endothelial hyaluran receptor-1), while intra-embryonic HE are negative for this marker (42). Confirming previous studies, the majority of VE-Cadherin^+^DLL4^+^ HE/EC in E8 murine yolk sac co-express LYVE1, whereas VE-Cadherin^+^DLL4^+^ HE/EC from the E9 embryo proper largely lack LYVE1 expression by flow cytometry (Figure 4A). This result is consistent with control E8.5-E9 embryo-derived cells (separated as yolk sac and embryo proper) analyzed in parallel with EBs in our sci-Plex data, in which *Lyve1* expression is predominantly detected in yolk sac HE/EC when compared to the embryo proper (Figure 4B, S3A).

**Figure 4:**
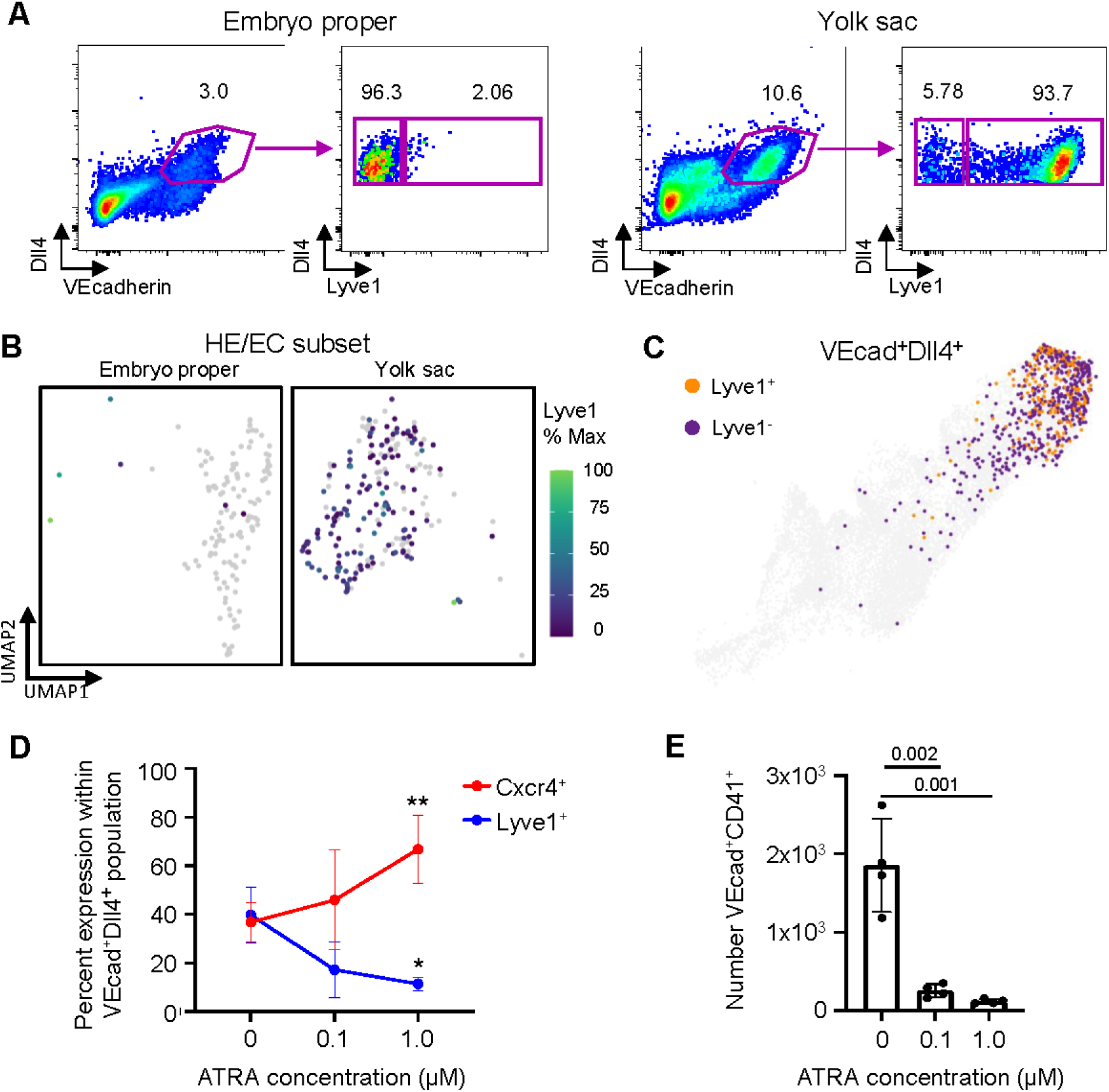
sci-Plex highlights production of intra- and extra-embryonic HE and a role for retinoic acid signaling. A: Expression of Lyve1 from embryo proper (left) and yolk sac (right) vascular tissues dissected from early E9 embryos. Cells were gated on Dll4^+^VECadherin^+^. Gates are drawn based on isotype controls. B: Gene expression heatmap for *Lyve1* expression for embryo proper (left) and yolk sac (right) tissues within the HE/EC subset of embryo dissections collected for sci-Plex. C: Gene expression of *Dll4^+^VECadherin^+^Lyve1^+/-^* within the HE/EC subset of EB sci-Plex data. D: Percent expression of Lyve1 (blue) and Cxcr4 (red) within the Dll4^+^VECadherin^+^ population across ATRA concentrations (see Figure S5B for associated dot plots). Asterisks designate significant difference compared to 0 μM. Error bars represent standard deviation. E: Number of VECadherin^+^CD41^+^ cells across ATRA concentrations (see Figure S5B for associated dot plots). P-values listed are comparison to 0 μM.

Consistent with these findings in vivo, we identified both yolk sac-like *Cdh5^+^Dll4^+^Lyve1^+^* EC/HE and *Cdh5^+^Dll4^+^Lyve1^-^* HE/EC in our EB dataset, a subset of which also expresses *Cxcr4*, consistent with intra-embryonic origin (6, 43) (Figure 4C and S3B). To verify generation of EC/HE subtypes from mESC in this protocol, we performed FACS isolation of EB cells to sort the VE-Cadherin^+^LYVE1^+^ and VE-Cadherin^+^LYVE1^-^ EC/HE (Figure S3C). The hematopoietic potential of sorted cell populations was assessed via co-culture on AGM-ECs, which we have previously shown can robustly support EHT and generation of multilineage hematopoietic progeny (6, 23). Both subsets generated CD45+ hematopoietic cells during co-culture (Figure S3D) and myeloid/erythroid colonies in methylcellulose assays (Figure S3E). B- and T-lymphoid potential, assessed by secondary OP9 and OP9-DLL4 stromal cells co-culture, respectively, demonstrated that LYVE1^-^ HE generated abundant CD4^+^CD8^+^ T-cells, while yolk-sac like LYVE1^+^ EC/HE did not (Figure S3F). However, the LYVE1^+^ population did generate B1a-like cells (AA4.1^+^CD19^+^B220^-/lo^; Figure S3G), an embryonic restricted B-cell subset predominantly emerging in the yolk sac (9). Taken together, these findings are consistent with the hematopoietic potential of their in vivo counterparts.

Although we did observe the natural progression of hematovascular markers, including Runx1, within the HE/EC subset (Figure S4A) and detect a minor population of *Cdh5^+^Dll4^+^Cxcr4^+^* EC/HE in our sci-Plex data (Figure S3B) suggestive of HSC-competent HE based on our prior studies in vivo (6, 41), several other notable genes that drive EHT in HSC-competent HE (egs *Gfi1*, *Hlf*, *Myb*), as reported by our lab and others (6, 44, 45), are distinctly lacking in our EB dataset (Figure S4B). This underscores the notion that acquisition of key transcriptional programs prior to undergoing EHT are necessary to endow HSC potential and highlights the need for future studies into driving expression of these critical programs to achieve HSC fate from mESC in vitro.

### sci-Plex highlights a role for retinoic acid signaling in HE fate

Retinoic acid (RA) signaling has previously been shown to inhibit yolk sac-like HE and promote intraembryonic AGM-like type HE from human PSC *in vitro,* and is required for development of long-term HSC in the AGM *in vivo* (46, 47). RA precursors are not present in serum-free media used in this protocol. However, a subset of EB-derived HE/EC do express RA receptors *Rarg* and *Rara*, indicating competence to respond to exogenous retinoic acid (Figure S5A). Hence, we tested whether addition of all trans retinoic acid (ATRA) to EB cultures on day 5 altered HE/EC cell fates. We found that while exogenous ATRA did not impact total cell numbers or generation of VE-Cadherin^+^DLL4^+^ arterial EC/HE (Figure S5B and C), it significantly enhanced frequency of VE-Cadherin^+^DLL4^+^ cells expressing CXCR4^+^ while reducing those expressing LYVE1^+^, in a dose-dependent manner (Figure 4D). ATRA also significantly reduced numbers of VE-cadherin^+^CD41^+^ cells (Figure 4E), suggesting inhibition of yolk sac-like hematopoiesis that predominates in the cultures at this stage. These results are consistent with huiPSC studies showing ATRA inhibited primitive hematopoiesis but improved definitive (47). However, we have not observed reproducible HSC activity from ATRA-treated mESC-derived EC/HE based on transplantation assays following AGM-EC co-culture (not shown), suggesting that additional exogenous factors necessary to promote efficient induction of HSCs from HE at this stage in the protocol remain to be determined in future studies.

## Discussion

In this study, we present several advances towards elucidating the complex, combinatorial signaling pathways involved in driving early cell fate decisions and manipulating those pathways to optimize in vitro hematopoietic differentiation of mESCs. First, we provide a proof of principle for utilizing sci-Plex scRNA-seq to dissect the complexities of dose and temporal integration of interacting signal pathways to determine cell fates. Next, we created a single cell atlas of early mesoderm patterning and various downstream lineages that can be used for discovery of additional cell fate choices. Finally, we used sci-Plex to optimize the production of intra-embryonic mesodermal fates, including LPM and HE, which serve as the precursors to HSC.

Here, we employed sci-Plex to decipher the combinatorial dose and time-dependent effects of an Activin/BMP4 gradient on mESC-derived mesoderm and hematopoietic lineages. High-throughput single cell methodologies such as sci-Plex are essential to dissect the complex nature of cell signaling pathways during early embryonic cell fate decisions, notably the subtle changes in timing and/or dosage of morphogens/cytokines which can have profound ramifications on downstream cell types. While sci-Plex has previously been applied to identify heterogeneity in drug response (22) or understand cell fate determinants in retinal organoids (48), here we utilized sci-Plex to deconvolute the complex, heterogeneous cellular landscape of an actively differentiating pluripotent stem cell population in response to combinations of morphogens. To this end, we were able to define the developmental trajectory of hematopoiesis accurately and confidently within our EBs, based on published markers and gene sets (Figure 1, 2 and Figure S2). This underscores the notion that sci-Plex allows for high throughput single-cell screening of a population with adequate resolution to distinguish subtle cell types and transitions in developmental states. Furthermore, it suggests that sci-Plex could have profound applications toward modeling both normal development and disease processes in hPSC and human organoid models. For example, sci-Plex could be employed to better understand the heterogeneity within the tumor microenvironment of various cancers, notably the interactions/signaling between cancer stem cells and the niche, in order to highlight novel therapeutic targets, and subsequently observe the cellular/transcriptional response to such perturbations.

The data generated in our study provides a robust single cell atlas that accurately recapitulates expected cell fates in early embryonic patterning in response to Activin/BMP4 gradients. Of note, our transcriptional analysis identified lineage pathways undertaken by mESCs differentiating primarily into various mesodermal lineages including cardiomyocytes, allantois, mesenchyme, and hematovascular cell types (Figure 1C and D). With each of these lineage pathways, there are key cell fate choices being made at the transcriptional level that are the primary drivers of subsequent differentiation, many of which remain poorly characterized. Here, we focused on defining some of those cell fate decisions important for generation of intra-embryonic HE, which serve as precursors to HSCs. However, this dataset provides a robust resource for analysis of other novel cell fate decisions during mesodermal patterning and differentiation resulting from combinatorial Activin/nodal and BMP4 signals.

We utilized sci-Plex to assess intra-embryonic vs extra-embryonic fates and optimize production of LPM, including HE, within our mESC differentiation protocol. Notably, we observed that higher Activin concentrations coupled with lower BMP4 favored IEC output and that LPM production peaked at day 5 of our protocol in 20 ng/mL Acitivin and 5 ng/mL BMP4 conditions (Figures 2D-E and 3A). Moreover, the transition from LPM to hematovascular fates, specifically HE, requires acquisition of endothelial programs (34, 39), which were inhibited by high Activin (Figure 3B-C), leading to a revised protocol with a lower Activin concertation after day 5. This optimization led to increased production of both yolk sac-like and intraembryonic-like HE, which we furthered defined through expression of LYVE1 and subsequent lineage assays (Figure 4 and FigureS3). Moreover, we observed these two HE populations are differentially responsive to RA signaling consistent with findings in human PSC (47).

Ultimately, while we were able to utilize sci-Plex to enhance production of HE, we did not observe long-term engraftment following transplantation into mice, suggesting there are additional barriers to efficient generation of HSC-competent HE. Of note, we observed minimal or absent expression of several key genes identified by our lab and others that define AGM arterial HE in vivo (6, 11, 45, 49). This may be, in part, due to the lack of key niche cell types and signaling pathways in the EBs that are required for HSC fate in vivo (50) and/or critical aspects of developmental timing required for maturation of HSC-competent HE (6). Thus, future studies are warranted to identify additional exogenous factors and critical temporal cues required for HSC competence that are not recapitulated in our current EB system. Furthermore, application of sci-Plex would enable high-throughput optimization of the signals and developmental timing that regulate HSC emergence in recent hiPSC platforms (12, 13), with the potential to improve both the efficiency of HSC generation and the engraftment properties of the resulting cells.

## Conclusions

Taken together, the results from this study underscore the importance of deciphering the complex, combinatorial signaling that occurs during early cell fate decisions and how that knowledge can be applied to optimize in vitro differentiation of pluripotent stem cells to a desired cell type or lineage. High-throughput tools, like sci-Plex utilized here, which allow researchers to measure transcriptional responses to highly multiplexed perturbations at single-cell resolution in a time-efficient and cost-effective manner will be a key aspect in any subsequent investigations. Thus, we present this study as a resource for the dataset generated on early mesodermal patterning, as well as a roadmap for the design of future studies to optimize specific cell fates in dynamic stem cell differentiation protocols.

## List of abbreviations

HSC: hematopoietic stem cell
mESC: murine embryonic stem cell
huiPSC: human induced pluripotent stem cell
EC: endothelial cell
HE: hemogenic endothelium
EHT: endothelial to hematopoietic transition
AGM: aorta-gonad-mesonephros region
PS: primitive streak
IEC: intra-embryonic cells
EEC: extra-embryonic cells
LPM: lateral plate mesoderm
EMP: erythromyeloid progenitor
AGM-EC: AGM-derived Akt-EC
EB: embryoid body
VEGF: vascular endothelial growth factor
BMP4: bone morphogenetic protein 4
ATRA: all-trans retinoic acid
SCF: stem cell factor
FLT3L: FMS-like tyrosine kinase 3 ligand
TPO: thrombopoietin
IL-3: interleukin-3
IL-7: interleukin-7
CFU: colony forming unit
CFU-E: CFU-Erythroid
CFU-GM: CFU-Granulocyte/Macrophage
CFU-GEMM: CFU-Granulocyte, Erythrocyte, Monocyte, Megakaryocyte
UMAP: Uniform Manifold Approximation
PGC: primitive germ cell
GSS: gene-set scores
LYVE1: lymphatic vessel endothelial hyaluran receptor-1

## Declarations

### Ethics approval

All animal studies were conducted in accordance with the NIH guidelines for humane treatment of animals and were approved by the Institutional Animal Care and Use Committee at the Fred Hutchinson Cancer Center.

### Availability of data and materials

All raw and processed sequencing data is publicly available through GEO (accession number GSE343689). All scripts for single cell transcriptomic analysis will be publicly available on the FHCC GitHub at https://github.com/FredHutch/mESC-SciPlex. Other datasets generated and/or analyzed during the current study are available from the corresponding author on reasonable request.

## Authors’ contributions

B.H., C.T., S.R.S., B.V., and A.M.H. conceived the study and designed experiments. B.V., A.M.H., S.R.S., S.D., B.H., T.D. and C.N. performed experiments and analyzed the data. B.V, A.M.H., B.H., S.R.S., R.W., and A.B.C. wrote and revised the manuscript. All authors reviewed and approved the final manuscript.

## Supplemental Figures

**Figure S1:**
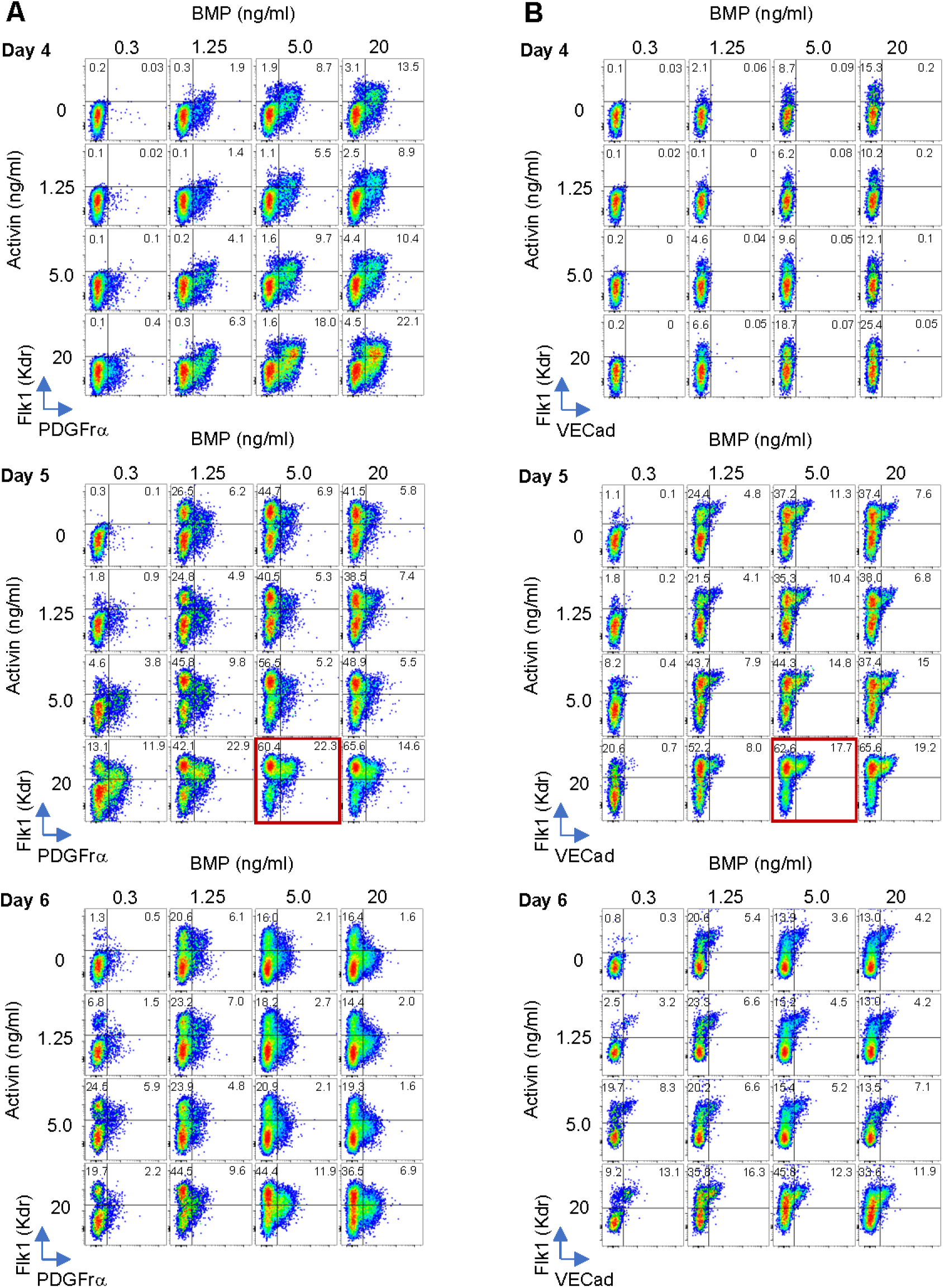
Phenotypic analysis of sci-Plex conditions. Expression of Flk1 (Kdr) vs Pdgfrα (A) or VECadherin (Cdh5) (B). Gates are drawn based off isotype controls. Red boxes indicate chosen conditions for optimal lateral plate mesoderm production and subsequent hematopoietic output.

**Figure S2:**
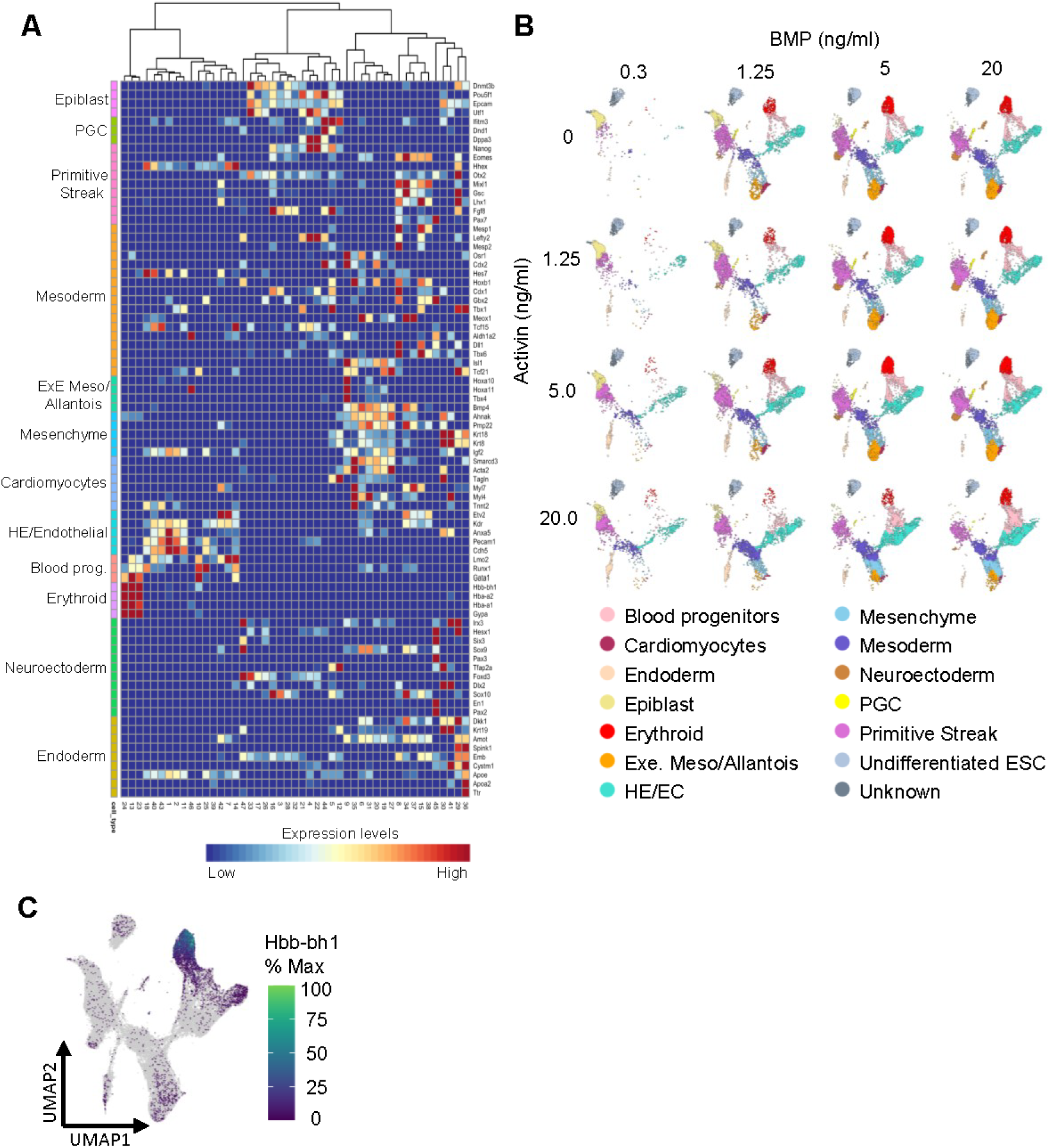
Cell type classification of sci-Plex dataset. A: Heatmap of diagnostic marker expression for cell type based on cluster. B: Cell type classification based on diagnostic markers subset for Activin/BMP conditions. C: Gene expression heatmap for primitive erythrocyte marker hemoglobin Z, beta-like embryonic chain (*Hbb-bh1*) within the full EB sci-Plex dataset.

**Figure S3:**
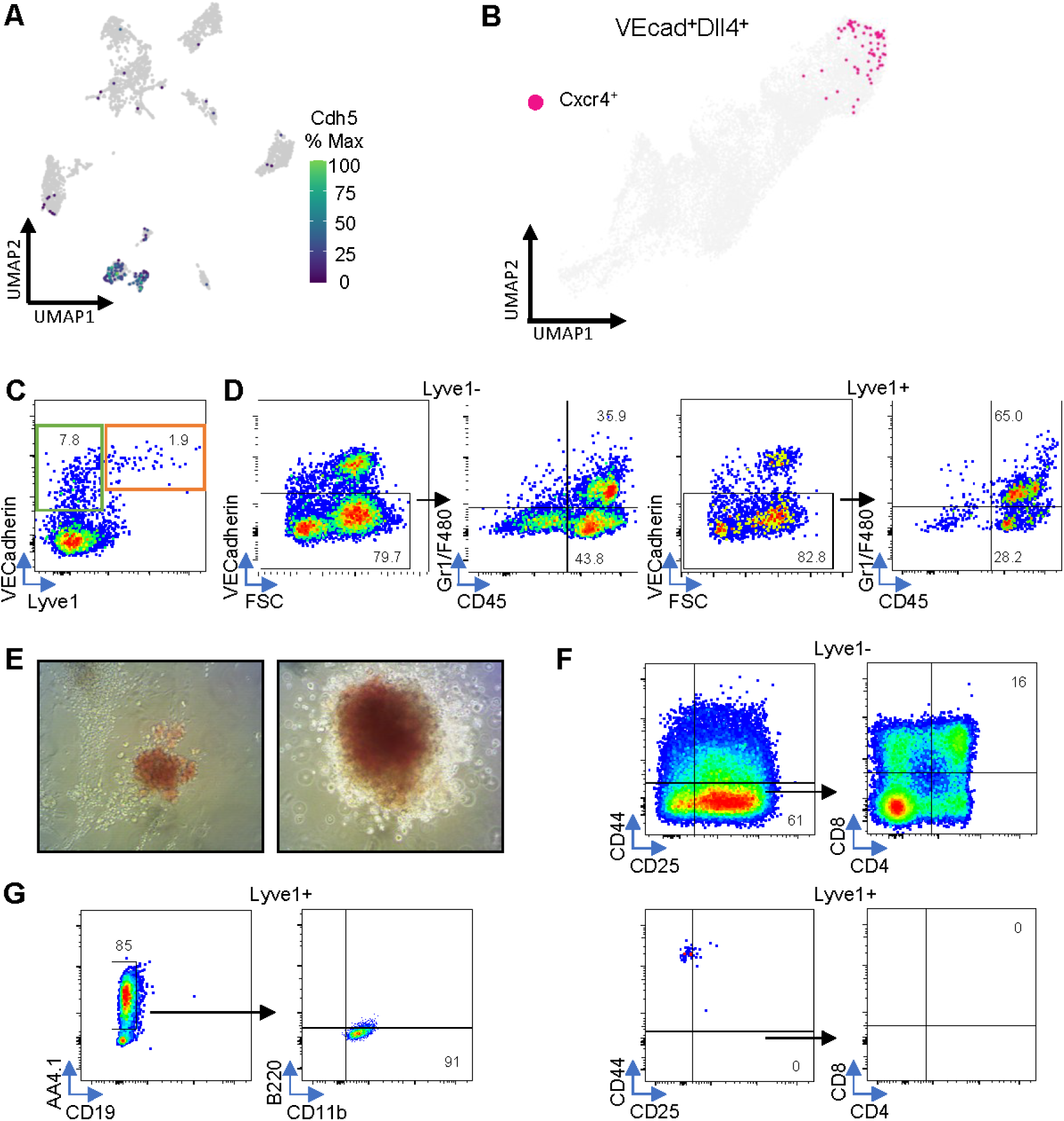
Functional assessment of hematopoietic activity for Lyve1^+/-^ populations generated from EB differentiation. A: Gene expression heatmap for VECadherin (*Cdh5*) within embryo dissection sci-Plex dataset. B: Gene expression of *Dll4^+^VECadherin^+^Cxcr4^+^* within the HE/EC subset of EB sci-Plex data. C: Expression of VECadherin and Lyve1 from EBs collected for sorting on Day 7 of differentiation. Green and orange boxes indicate the sorted populations. D: Phenotypic expression of Lyve1^-^ (left) and Lyve1^+^ (right) cells sorted from (C) for the pan-hematopoietic marker CD45 and myeloid markers Gr1 and F480 within the VECadherin^-^ population of cells after 13 days of co-culture. E: Images of CFU-E (left) and CFU-GEMM (right) colonies generated in CFU assay for cells collected from (D). F: Phenotypic expression of Tcell markers CD44^-^CD25^+^CD8^+^CD4^+^ for Lyve1^-^ (top) and Lyve1^+^ (bottom) cells collected from (D) and subjected to co-culture on OP9-Dll4 stromal cells. G: Phenotypic expression of Bcell markers CD19^-^AA4.1^+^B220^-^CD11b^+^ for Lyve1^+^ cells collected from (D) and subjected to co-culture on OP9 stromal cells. For all dot plots in C-G, gates are drawn based off isotype controls.

**Figure S4:**
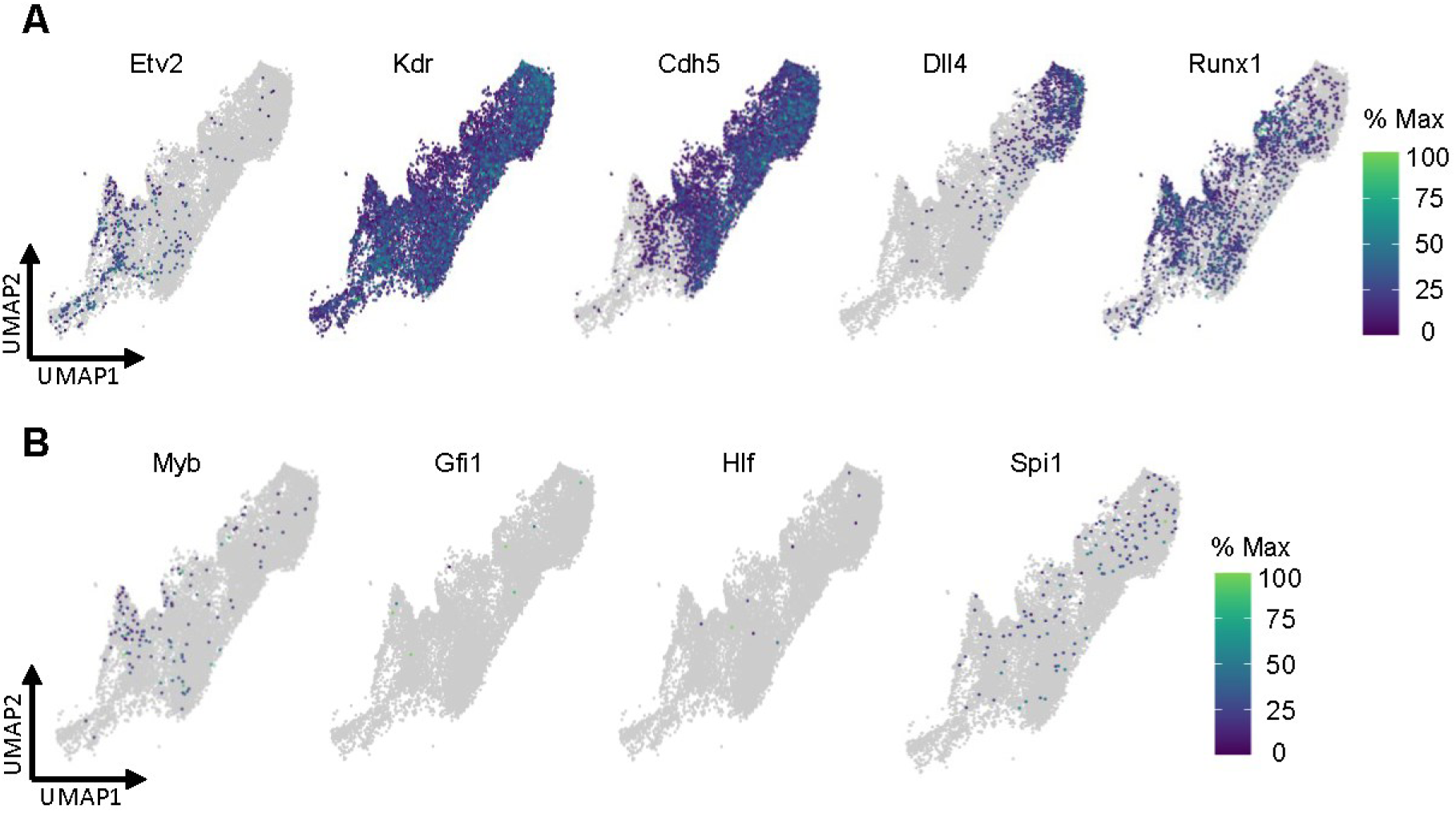
Expression of key hematopoietic development markers from sci-Plex data. Gene expression heatmaps for hematovascular (A) and EHT (B) markers within the HE/EC subset of EB sci-Plex data.

**Figure S5:**
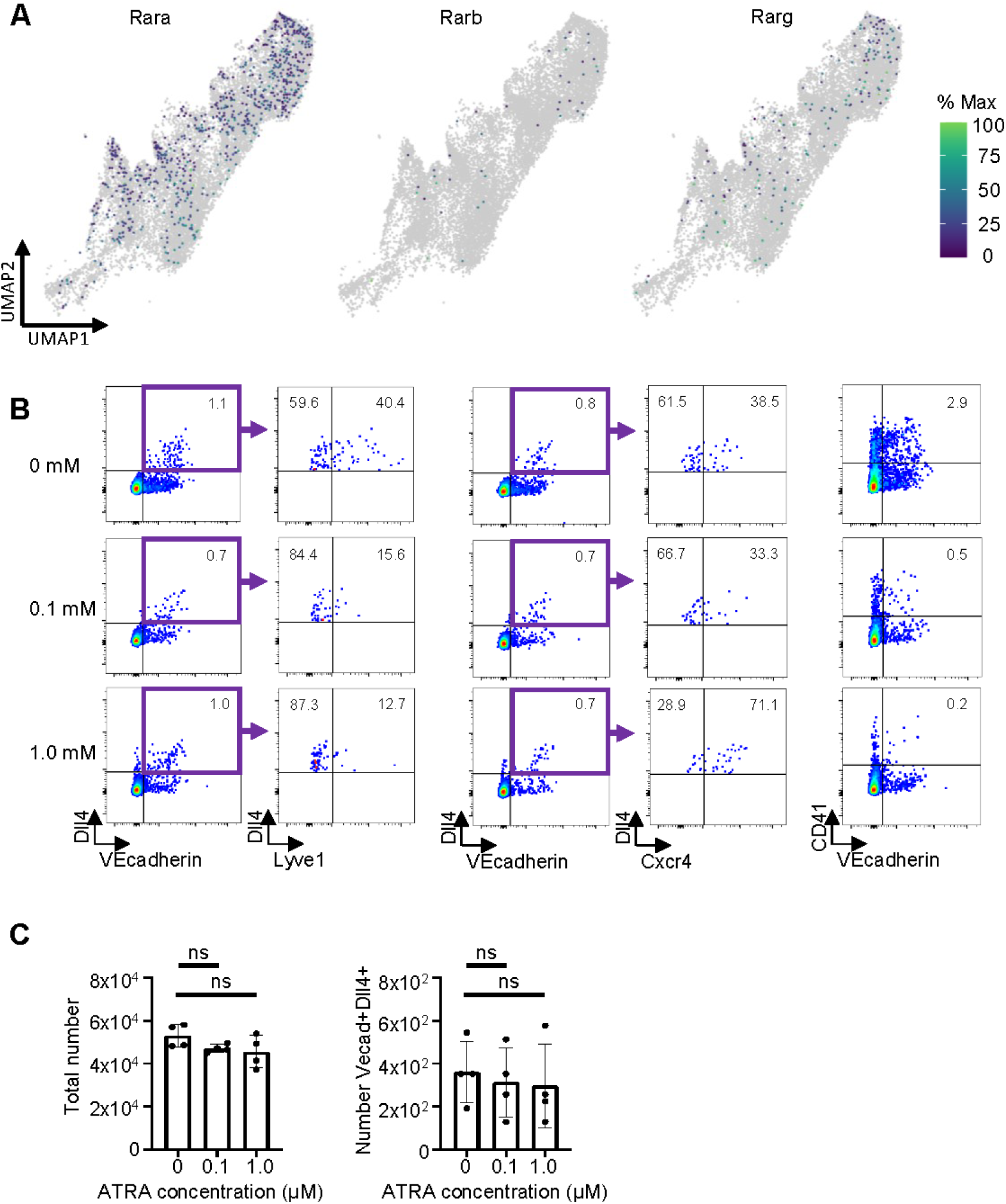
Retinoic acid signaling influences HE fate in EB differentiation protocol. A: Gene expression heatmaps retinoic acid receptors within the HE/EC subset of EB sci-Plex data. B: Phenotypic expression of Lyve1 (left) and Cxcr4 (middle) within the Dll4^+^VECadherin^+^ population, and CD41 x VECadherin (right). Gates are drawn based off isotype controls. C: Total (left) or VECadherin^+^Dll4^+^ number of cells across ATRA concentrations. Error bars represent standard deviation. Comparisons to 0 μM were non-significant, as listed.

